# Mechanisms of Differential Signal Transduction by IFNLR1 Variants

**DOI:** 10.1101/2025.10.03.677101

**Authors:** Laura A. Novotny, Carla Martinez-Morant, Stephen A. Duncan, Paula Traktman, Monika Gooz, Eric G. Meissner

## Abstract

**Background:** Lambda interferons bind the interferon lambda receptor-1 (IFNLR1) and IL10RB heterodimer to induce interferon stimulated genes (ISGs) and impart antiviral immunity. We previously showed that protein variants derived from distinct *IFNLR1* splice isoforms uniquely influence gene expression and HBV replication in stem cell-derived hepatocytes (iHeps). Here, we evaluated the molecular mechanisms of signal transduction by full-length canonical IFNLR1 (variant 1) and a non-canonical variant missing a portion of the Box1 and Box2 JAK1-interacting motifs (variant 2).

**Methods:** We used HEK293T cells, wild-type (WT), and *IFNLR1* knock-out (KO) iHeps that stably express doxycycline-inducible, FLAG-tagged IFNLR1 variants to evaluate function. Cellular responses to IFNL were measured using the Duolink proximity ligand assay, ImageStream flow cytometry, western blotting of JAK-STAT proteins, susceptibility to JAK1 and TYK2-specific inhibitors, and gene expression profiling.

**Results:** While each IFNLR1 variant colocalized with IL10RB after IFNL treatment, variant 1 was more rapidly and extensively internalized than variant 2. In WT iHeps with intact endogenous *IFNLR1*, expression of variant 2 enabled higher JAK1 and TYK2 phosphorylation than variant 1, yet contrarily, variant 1 enabled greater STAT1 and STAT2 phosphorylation that resulted in broader and higher expression of ISGs. Select ISGs exhibited differential constitutive expression in IFNL-untreated variant-expressing iHeps but were IFNL-inducible through both variants. In iHeps expressing variant 1, WT-iHeps were more resistant than KO-iHeps to TYK2-inhibition of antiviral ISG expression yet conversely were more susceptible to TYK2 inhibition of proinflammatory ISG expression, suggesting noncanonical variants made from endogenous *IFNLR1* transcripts influence the TYK2-dependence of IFNL signaling.

**Conclusions:** IFNLR1 variants promote differential utilization of signaling mediators to influence IFNL-induced gene expression patterns, indicating a putative role in pathway regulation.

## Introduction

Lambda interferons (IFNL) direct cellular immune responses to pathogens encountered at mucosal surfaces, including the liver, respiratory, and gastrointestinal tracts^1–14^. Soluble IFNLs bind the interferon lambda receptor 1 (IFNLR1) which heterodimerizes with the interleukin 10 receptor beta (IL10RB) to promote transphosphorylation of JAK1 bound to IFNLR1 and TYK2 bound to IL10RB^15–18^. Subsequent phosphorylation of docking sites on the IFNLR1 intracellular domain results in recruitment and phosphorylation of STAT proteins that promote induction of antiviral interferon stimulated genes (ISGs), similar to type-I IFN signaling; however, type-I IFNs also induce proinflammatory ISGs while IFNLs do not^3,19–21^. This may relate in part to the higher relative expression levels of the interferon alpha receptor 1 and 2 (IFNAR1/2) subunits on most nucleated cells^6,17,22–30^, as IFNLR1 overexpression enables increased IFNL-induced expression of antiviral ISGs and *de novo* expression of proinflammatory ISGs^31–35^. Low and restricted IFNLR1 expression on epithelial cells at immune-tolerant locations may thus serve a regulatory function to permit antiviral responses while restricting harmful inflammation^36–38^.

To limit adverse sequelae of excessive signaling, type-I IFNs trigger IFNAR1/2 internalization/degradation and induce ISG proteins that bind and inhibit IFNAR1/2; further, cells express truncated IFNAR2 variants from *IFNAR2* splice isoforms that inhibit type-I signaling^31,39–50^. In comparison, IFNLR1 is less impacted by the proteins that negatively regulate IFNAR1/2^51–53^, but other mechanisms cells use to regulate IFNL signaling are not fully established. Through unclear mechanisms, persons with genetic TYK2-deficiency and those treated with TYK2 inhibitors retain IFNL but not type-I IFN signaling capacity^54–56^, indicating that TYK2 bound to IL10RB is not an absolute requirement for IFNL signaling. Furthermore, work with synthetic IFNLR1 receptors harboring the extracellular domain of the erythropoietin receptor suggested that TYK2 is required for signaling through the IFNLR1/IL10RB heterodimer^57–59^. In contrast, synthetic IFNLR1 homodimers, each bound to JAK1, propagate TYK2-independent signaling using STAT proteins, suggesting noncanonical receptor signaling complexes that lack or do not require TYK2 and/or IL10RB could participate in IFNL signal transduction^57–59^. Work with synthetic, recombinant receptors also identified that differences in the cytoplasmic JAK1 binding motifs between IFNAR2 and IFNLR1 contribute to their differential capacity to transduce signal^59^. Gaining further insight into IFNL signaling regulation through study of native IFNLR1 protein has been difficult due to the intrinsically low IFNLR1 expression on cells that has challenged its detection and study^32,60–62^.

Intriguingly, RNA sequencing of primary cells identified not only full-length canonical *IFNLR1*, but also noncanonical, truncated *IFNLR1* splice isoforms that encode for proteins that are conceptually similar to IFNAR2 variants, but whose function are unclear^15,22,32,63,64^. Noncanonical *IFNLR1* transcripts encode for a variant that lacks part of the cytoplasmic Box1 and Box2 domains that bind and stabilize JAK1 (variant 2) and for a secreted variant that lacks a transmembrane domain (variant 3)^32,63,64^.

Recombinant IFNLR1 variant 3 was shown to bind and sequester IFNLs at the cell surface to limit ISG induction^22,65^, but little is known about variant 2. Higher relative expression of *IFNLR1* isoform 1 correlates with the capacity of IFNLs to induce ISGs in cells^22,32,63–66^, suggesting noncanonical IFNLR1 variants could function to regulate canonical IFNLR1; however, the mechanisms of this potential regulation are not established, and protein expression of noncanonical IFNLR1 variants has not been demonstrated in primary cells^67^.

In our prior work evaluating IFNLR1 variant function in HEK293T cells and stem cell-derived hepatocytes (iHeps), we identified that variant 1 enabled higher IFNL-induced antiviral ISG expression, *de novo* proinflammatory ISG expression, and enhanced inhibition of hepatitis B virus (HBV) replication^33–35^. In contrast, variant 2 enabled a partial increase in IFNL-induced antiviral ISG expression but did not support proinflammatory ISG induction and minimally impacted HBV^33–35^. Here, we evaluated the molecular mechanisms of signal transduction by IFNLR1 variants 1 and 2 to further explore their potential role in IFNL signaling regulation.

## Materials and Methods

### Cell culture

HEK293T clones with stable expression of doxycycline (dox)-inducible FLAG-tagged *IFNLR1* isoform constructs (referred to as FLAG-Iso1, FLAG-Iso2) or an empty vector control (EV) were derived and maintained as previously described^33^. K3 induced pluripotent stem cells (iPSCs) with intact endogenous *IFNLR1* (wild type; WT) or CRISPR-Cas9-mediated knock-out of *IFNLR1* expression (*IFNLR1*-KO) and stable expression of dox-inducible FLAG-Iso1, FLAG-Iso2, or EV constructs were derived as previously described and differentiated to iHeps over 20 days with alternating normoxic and hypoxic conditions^34,35,68,69^. Culture medium was supplemented with 1µg/ml puromycin and 10ng/ml doxycycline hyclate (Millipore-Sigma) during differentiation to prevent construct silencing^34,35,70,71^. Differentiated iHeps were maintained in hepatocyte culture medium with 20ng/ml oncostatin and 1µg/ml puromycin at 37°C and 5% CO_2_.

### Proximity ligation assay (PLA)

HEK293T FLAG-Iso1, FLAG-Iso2, and EV cells were seeded at 50% confluency on collagen I-coated 18-well chamber slides (Ibidi), incubated 24h without dox, then stimulated +/-IFNL3 (100ng/ml, R&D Systems) for 15min. Cells were fixed (4% paraformaldehyde), blocked for 1h (3% BSA, 0.1% TritonX-100 in PBS), then incubated with murine anti-FLAG (clone M2; Sigma-Aldrich) and rabbit anti-IL10RB (PA5-112888; ThermoFisher) overnight at 4°C. PLA was performed using the Duolink In Situ PLA mouse/rabbit kit (Millipore-Sigma) according to the manufacturer’s instructions. Controls included cells incubated with individual antibodies or no antibody. Images were captured on a Zeiss 510 confocal scanning microscope and rendered with Zeiss Zen software. Quantitation of the number of puncta per cell was performed on 3 representative images per condition using Duolink ImageTool software (Millipore-Sigma).

### Imaging flow cytometry

HEK293T FLAG-Iso1 and FLAG-Iso2 cells were dox-induced (100 ng/ml, 24h), stained with Live-or-Dye 615/740 viability stain (32015; Biotium) for 15min at 4°C, then washed by centrifugation with cold DPBS (2% FBS), then stained with rat anti-FLAG conjugated to Alexa Fluor 488 (637317; BioLegend) for 45min on ice. Controls included singly stained cells to establish live cell and FLAG-specific gates. After washing, cells were incubated +/-IFNL3 (100ng/ml) at 37°C for 0, 5, 15, or 30min, washed, then fixed for 15min on ice (4% paraformaldehyde). After two additional washes, cells were analyzed using a Cytek Amnis ImageStream Mk imaging flow cytometer with collection of 5,000 live events. Images were acquired in brightfield and fluorescent channels with a membrane mask generated on the cell periphery of the brightfield image using Amnis IDEAS 6.2 software. Mean fluorescence intensity within the mask at each timepoint was determined and normalized to values obtained at the zero-time point.

### Expression of endogenous *IFNLR1* isoform and FLAG-Iso transcripts

Expression of endogenous *IFNLR1* and FLAG-Iso transcripts was evaluated in iHeps treated +/-dox (100ng/ml) for 24h. Extracted RNA (Qiagen RNeasy) was quantified (Nanodrop) prior to cDNA synthesis (High-Capacity cDNA Reverse Transcription kit, Applied Biosystems) and quantitative reverse transcriptase-polymerase chain reaction (qRT-PCR) using commercial TaqMan (**Supplemental Table 1**) and custom primer-probes (for FLAG-Iso constructs as in^34^) with Fast Advanced Master Mix (Applied Biosystems). Assays were performed in biological and technical replicates with expression calculated relative to *GAPDH*.

### Western blotting

WT- and *IFNLR1*-KO-iHeps harboring FLAG-Iso1, FLAG-Iso2, and EV constructs were +/-dox-induced (100ng/ml, 24h) then mock, IFNL3 (100ng/ml), or IFNA2 (100ng/ml, PBL Assay Science) stimulated at 37°C for 15min, then collected and lysed on ice in RIPA buffer (ThermoFisher) supplemented with protease and phosphatase inhibitors (Pierce). Clarified protein (20µg) was separated on 10% Criterion TGX gels (BioRad) in SDS running buffer, transferred to nitrocellulose membranes (BioRad), then blocked in tris-buffered saline with 1% Tween20 (TTBS) and 5% Blotto (Santa Cruz Biotechnology). Membranes were incubated with primary antibodies diluted in TTBS plus 5% BSA or 5% Blotto overnight at 4°C, then probed with goat anti-rabbit IgG-HRP (R&D Systems) or goat anti-mouse IgG-HRP (Millipore-Sigma) in TTBS plus 5% Blotto for 1h at room temperature. Blots were developed with Clarity Western ECL reagent (BioRad), and images were captured on a BioRad ChemiDoc system. After detection of phosphorylated signaling mediators, membranes were incubated with Restore Plus Western Blot Stripping buffer (ThermoFisher) for 30min at room temperature prior to washing, blocking, and incubation with antibodies to total proteins. Rabbit monoclonal antibodies to pJAK1 Y1034/1035 (74129), pTYK2 Y1054/1055 (68790), pSTAT1 Y701 (9167), pSTAT2 Y690 (88410), JAK1 (3344), TYK2 (14193), STAT1 (14994), and STAT2 (72604) were from Cell Signaling Technologies. FLAG was detected with murine monoclonal anti-FLAG clone M2 (F1804, Millipore-Sigma) and equivalent protein per lane was evaluated with polyclonal rabbit anti-GAPDH (PAB932HU02, Cloud-Clone Corp). Integrated band intensity was quantitated for phosphorylated and total protein using ImageJ.JS software with a background correction performed within each lane. To account for potential differences in the available pool of each protein and enable comparisons between signaling mediators, the ratio of phosphorylated to total protein is shown, converted to percentage.

### Flow cytometry

Dox-uninduced HEK293T FLAG-Iso1 and FLAG-Iso2 cells were incubated with Live-or-Dye viability stain in PBS (15min), then treated +/-IFNL3 (100ng/ml, 15min) and collected by pipetting with fixation in 4% paraformaldehyde. Cells were washed twice with Dulbecco’s PBS plus 2% FBS with centrifugation at 500*xg*, permeabilized on ice with cold methanol (15min), then washed twice and incubated with murine monoclonal anti-pSTAT1 pY701 conjugated to PE (612564; BD) for 45min at room temperature. Cells were washed twice, then 20,000 live events were collected on a Millipore Guava 8HT flow cytometer with data analysis using FlowJo software (FlowJo, LLC).

### Susceptibility of IFNLR1 variants to JAK1 and TYK2 inhibition

Variant-expressing HEK293T cells and WT iHeps treated +/-dox (100ng/ml) and *IFNLR1*-KO iHeps treated + dox (100ng/ml dox) for 22h were incubated in medium +/-0.01-1.0µM upadacitinib (a JAK1 inhibitor, Tocris) or 0.01-1.0µM deucravacitinib (a TYK2 inhibitor, TargetMol) for 2h prior to stimulation +/- IFNL3 (100ng/ml, 24h) in the continued presence of inhibitor. RNA extraction and gene expression analyses were performed as above by qRT-PCR with percent expression calculated relative to similarly treated cells incubated without inhibitor.

### RNA sequencing

RNA sequencing was performed on variant-expressing WT and *IFNLR1*-KO iHeps and EV-controls^34,35^. Dox-uninduced WT-iHeps, dox-induced (100 ng/ml, 24h) *IFNLR1*-KO iHeps, and similarly treated respective EV controls were stimulated +/-IFNL3 (100ng/ml, 24h) before RNA extraction (Qiagen RNeasy kit). RNA sample purity and integrity were confirmed on an Agilent 5400 before library preparation and sequencing by Novogene (sample RNA Integrity Number (RIN) ranged between 8.8-9.9). Analysis of FASTQ files was performed using Partek Genomics Suite (v12.3.1) with read alignment using the STAR aligner, annotation to the human genome (vHg38), and normalization using the counts per million method. Differentially expressed genes (DEGs) between groups were identified using Deseq2 (R) with statistical significance achieved at p-value (≤ 0.05), fold change (-2,2), and false discovery rate (FDR; ≤ 0.05). Pathway Enrichment Analysis was performed on DEGs using Enrichr; pathways with an FDR <0.01 and at least 10 genes from the dataset represented in the specific pathway are reported. Heatmaps of DEGs were generated using GraphPad Prism (v10).

### Statistical analyses

Statistical analyses were performed using GraphPad Prism (v10) with data presented as mean ± standard error and statistical significance set at *p*≤ 0.05.

## Results

### IFNLR1 variants 1 and 2 dimerize with IL10RB after IFNL3 exposure

Relative to full-length canonical variant 1 (Iso1), IFNLR1 variant 2 (Iso2) lacks 29 amino acids in the cytoplasmic domain that encompass part of a Box1 motif that provides most of the energetics to support JAK1 binding^64,72^, including the last 2 amino acids of the critical PxxLxF motif^73^, and the majority of a Box2 motif that stabilizes the IFNLR1-JAK1 interaction^72^ (**Fig. 1**). To evaluate how the missing Box domain elements affect signaling, we first asked whether each variant could colocalize with IL10RB after IFNL3 treatment. Because endogenous IFNLR1 is difficult to visualize^32,60–62^, we utilized HEK293T cells with stable dox-inducible expression of FLAG-tagged IFNLR1 variants^33^ and performed a proximity ligation assay to examine FLAG and IL10RB colocalization. We used dox-uninduced cells, which express some construct transcript from a leaky tet-on promoter and which fully sensitize cells to the maximal effects of IFNL3 treatment^33^, in order to minimize the potential for off-target effects related to protein overexpression, which could alter the relative stoichiometry of signaling molecules. Neither variant associated with IL10RB in untreated cells, and no background signal was detected in IFNL3-treated EV cells (**Fig. 2A**). In IFNL3-treated cells expressing FLAG-Iso1 or FLAG-Iso2, we observed numerous puncta (**Fig. 2A**) with comparable levels per cell for each variant (**Fig. 2B**). These data indicated that each variant could form complexes with IL10RB upon ligand engagement. This likely relates to their identical extracellular domains that enable the critical stem-stem interactions with IL10RB that stabilize the ternary complex with ligand^73^. Moreover, these data and our prior work^33–35^ confirmed *IFNLR1* isoform 2, identified in primary cells as an alternatively spliced transcript but not yet shown to be translated to protein *in vivo*, encodes for a functional protein than engages with IFNLs and IL10RB to transduce signal.

**Figure 1.**
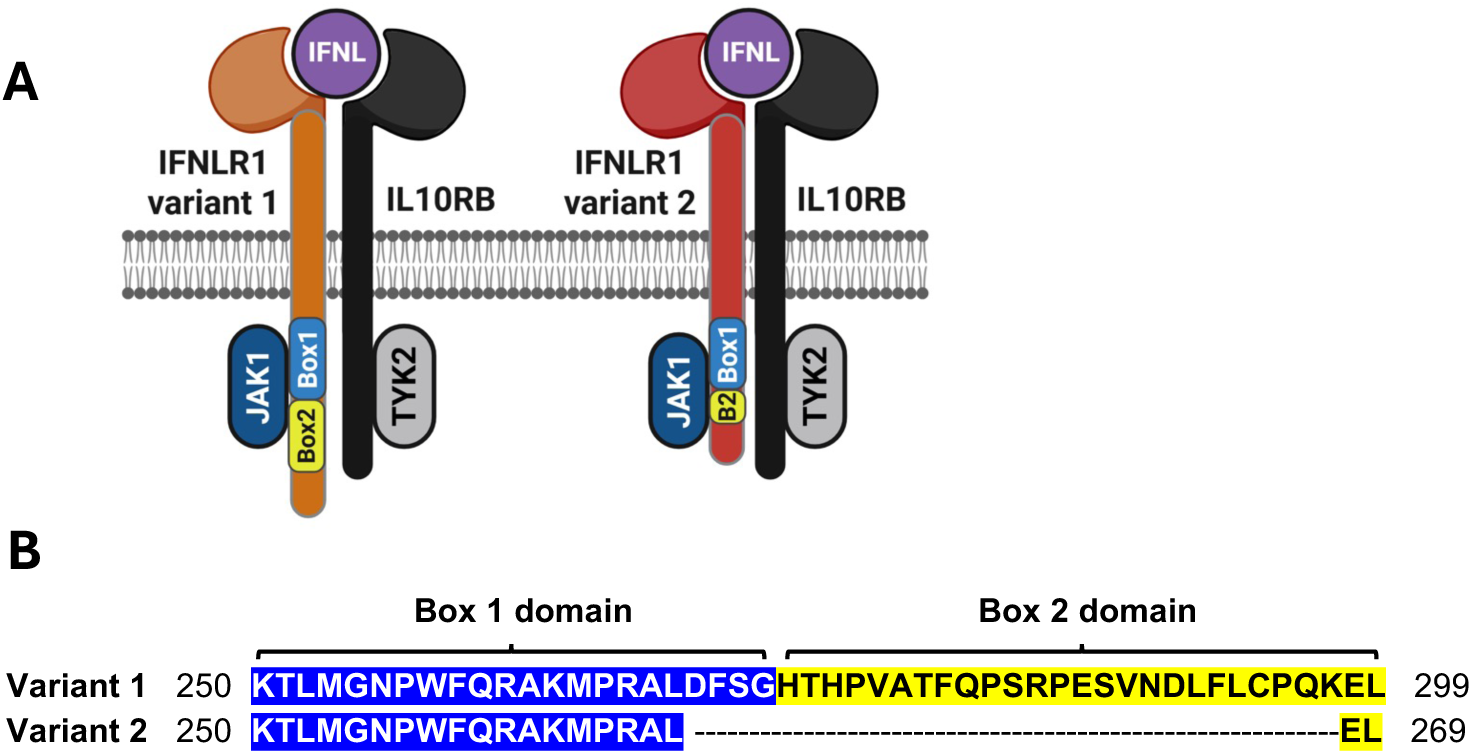
Schematic of IFNLR1 variants 1 and 2. **(A)** Cartoon depicting IFNL bound to each IFNLR1 variant and complexed with IL10RB with proximal kinases JAK1 and TYK2 engaged. Note the truncation in the Box1 and Box2 JAK1 binding motifs within IFNLR1 variant 2. **(B)** Alignment of the JAK1 binding motifs for IFNLR1 variant 1 (Q8IU57; top sequence) and IFNLR1 variant 2 (Q8IU57-2; bottom sequence). Box 1 (blue highlight) and Box 2 (yellow highlight) domains are indicated^72^ and reveal the 29 amino acid truncation within the cytoplasmic domain of IFNLR1 variant 2 that eliminates the C-terminal residues of Box 1 and all but two residues of Box 2. Created in BioRender. Novotny, L. (2025) https://BioRender.com/24aqcgc.

**Figure 2.**
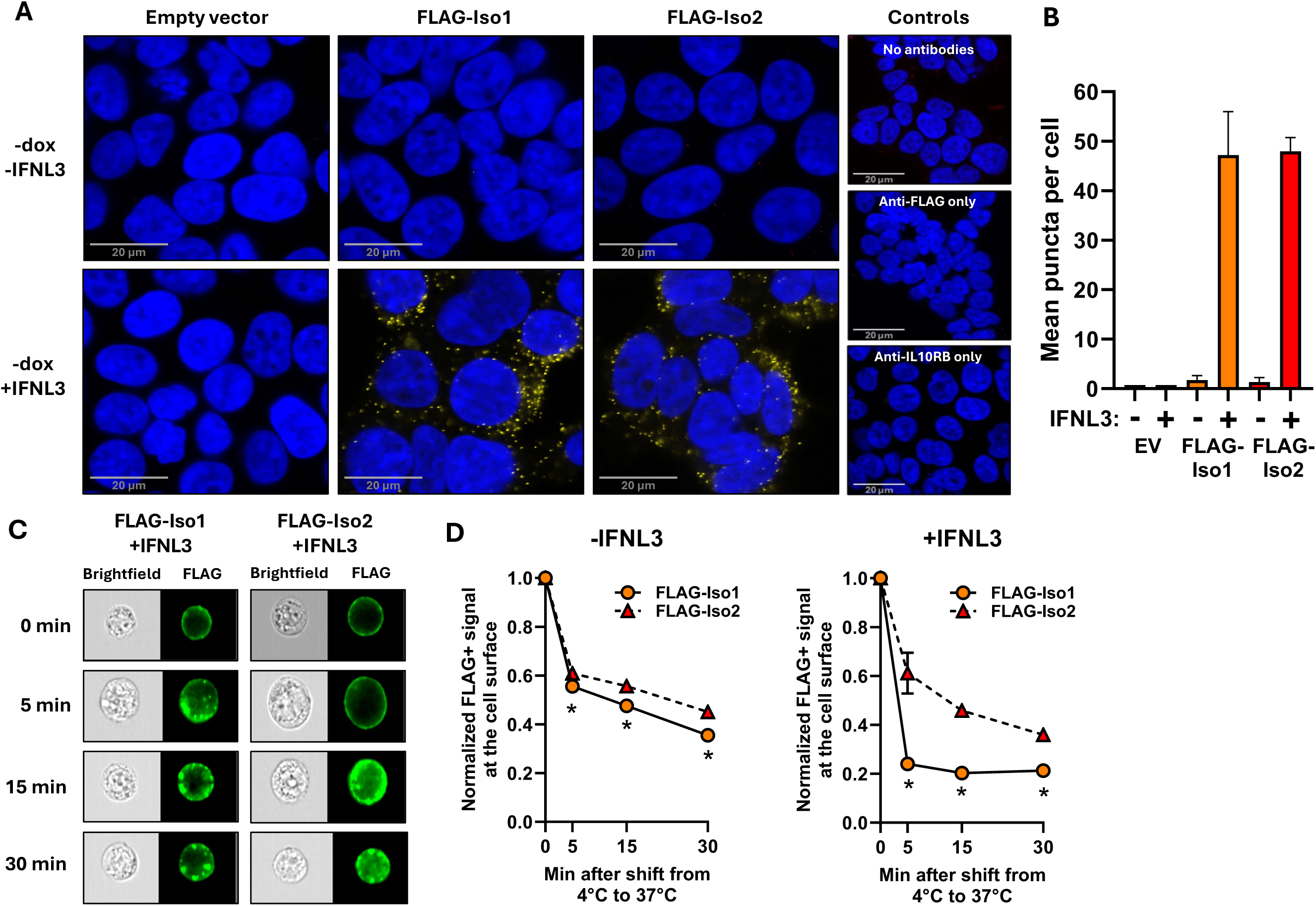
IFNLR1 variants 1 and 2 bind IFNL3 and complex with IL10RB but have differing internalization kinetics. **(A)** Duolink PLA was performed to examine interactions of FLAG-Iso1 and FLAG-Iso2 with IL10RB after IFNL3 stimulation (15min). Yellow puncta represent complex formation; nuclei were counterstained with DAPI (blue) and imaged by scanning confocal microscopy. Scale bar, 20µm. **(B)** Quantitation of mean puncta per cell showing comparable ability of FLAG-Iso1 and -Iso2 to form ligand-induced complexes with IL10RB. Three images per condition were analyzed, mean ± SEM shown. n=43 cells per image. (**C**) Representative brightfield and fluorescent images of dox-induced FLAG-Iso1 and –Iso2 cells pre-incubated with anti-FLAG antibody at 4°C, then shifted to 37°C in the presence or absence of IFNL3 showing redistribution of FLAG-specific fluorescent signal (green) over time. Images selected at each time point represent cells with comparable peripheral FLAG-specific fluorescent signal that corresponded with the mean observed for the total population. **(D)** Quantitation of FLAG-specific signal at the cell surface, normalized to the respective time point zero. n= 5000 live events, mean ± SEM shown. *p≤ 0.05, comparing FLAG-Iso1 to –Iso2 at each timepoint.

### FLAG-Iso1 is more rapidly internalized than FLAG-Iso2

We next used high-throughput imaging flow cytometry to compare the receptor internalization kinetics of each variant by quantifying surface to cytoplasmic FLAG levels in untreated and IFNL3-treated conditions. Attempts to optimize laser intensity and imaging sensitivity to enable fluorescence detection in dox-uninduced cells were unsuccessful; therefore, we performed this assay in dox-induced HEK293T cells. Both FLAG-Iso1 and FLAG-Iso2 populations had comparable surface labelling with anti-FLAG antibody at 4°C (mean fluorescent intensity of 5.0x10^4^ and 4.9x10^4^, respectively). Upon transition to 37°C to allow internalization, we detected redistribution of fluorescence from the cell periphery to the intracellular region within punctate structures suggestive of endosomes (**Fig. 2C**). Over time, FLAG-Iso1 was more rapidly and extensively internalized than FLAG-Iso2 in both untreated and IFNL3-treated conditions (**Fig. 2D**), suggesting the missing cytoplasmic region of variant 2 slowed its capacity to be internalized.

### IFNLR1 variants differentially support phosphorylation of JAK-STAT proteins

Due to the missing elements of the Box1 and Box2 JAK1-binding motifs (**Fig. 1**) and based on prior structural work^64,72^, we hypothesized that variant 2 may have reduced JAK1 affinity, which could impair its capacity to promote JAK1, TYK2, and/or STAT1/2 phosphorylation and thus influence downstream signaling. To test this, we evaluated IFNL-induced phosphorylation of JAK-STAT proteins in WT and *IFNLR1*-KO iHeps expressing each variant. We evaluated both dox-uninduced and dox-induced conditions, but prioritized the quantitative analysis of dox-uninduced WT iHeps, which express low levels of construct that are similar to endogenous transcript levels (**Supplemental Fig. 1**), to mitigate the risk of off-target effects related to protein overexpression. We also evaluated *IFNLR1*-KO iHeps, however these cells required dox-induction and higher construct expression levels to have an observable phenotype^33–35^. We compared phosphorylated to total protein for each analyte and observed no notable differences in total protein levels between samples.

While both variants in dox-uninduced WT-iHeps enabled increased pJAK1 and pTYK2 relative to EV control iHeps, FLAG-Iso2 unexpectedly supported higher levels of pJAK1 and pTYK2 than FLAG-Iso1, yet lower levels of pSTAT1 and pSTAT2 (**Fig. 3A&C**; dox-induced quantitative data is shown in **Supplemental Figure 2A**) despite both variants containing all of the cytoplasmic domain tyrosines implicated in STAT1/STAT2 binding and phosphorylation^31,37,74,75^. In dox-induced *IFNLR1*-KO iHeps, FLAG-Iso1 enabled higher IFNL3-induced pJAK1, pTYK2, and pSTAT2 than FLAG-Iso2 whereas pSTAT1 levels were similar (**Fig. 3B&D**; dox-uninduced quantitative data shown in **Supplemental Figure 2B**). These differences in phosphorylated signaling mediators were specific to IFNL signaling, as neither variant had a notable effect on IFNA2 signaling in dox-uninduced WT- and KO-iHeps, with the exception of pTYK2 which was higher in IFNA2-treated KO-Iso1 and KO-Iso2 relative to KO-EV iHeps (**Fig. 3A-B, Supplemental Figure 2C-F**). We did observe potential off-target dampening of type-I IFN signaling in dox-induced *IFNLR1*-KO iHeps (**Fig. 3A-B, Supplemental Figure 2C-F**). Consistent with our pSTAT1 observations in dox-uninduced variant-expressing WT-iHeps (**Fig. 3A&C)**, dox-uninduced HEK293T FLAG-Iso1 cells had higher IFNL3-induced pSTAT1 than FLAG-Iso2 cells by flow cytometry (**Supplemental Figure 3**). These data confirmed IFNLR1 variant 2 can independently facilitate IFNL-induced recruitment and phosphorylation of JAK1 and TYK2, correlating with its capacity to complex with IL10RB and support IFNL-induced antiviral ISG expression (**Fig. 2** and ^33–35^), however the missing cytoplasmic Box domain elements restricted STAT1/2 phosphorylation relative to full-length variant 1, most notably in WT iHeps with intact endogenous *IFNLR1*.

**Figure 3.**
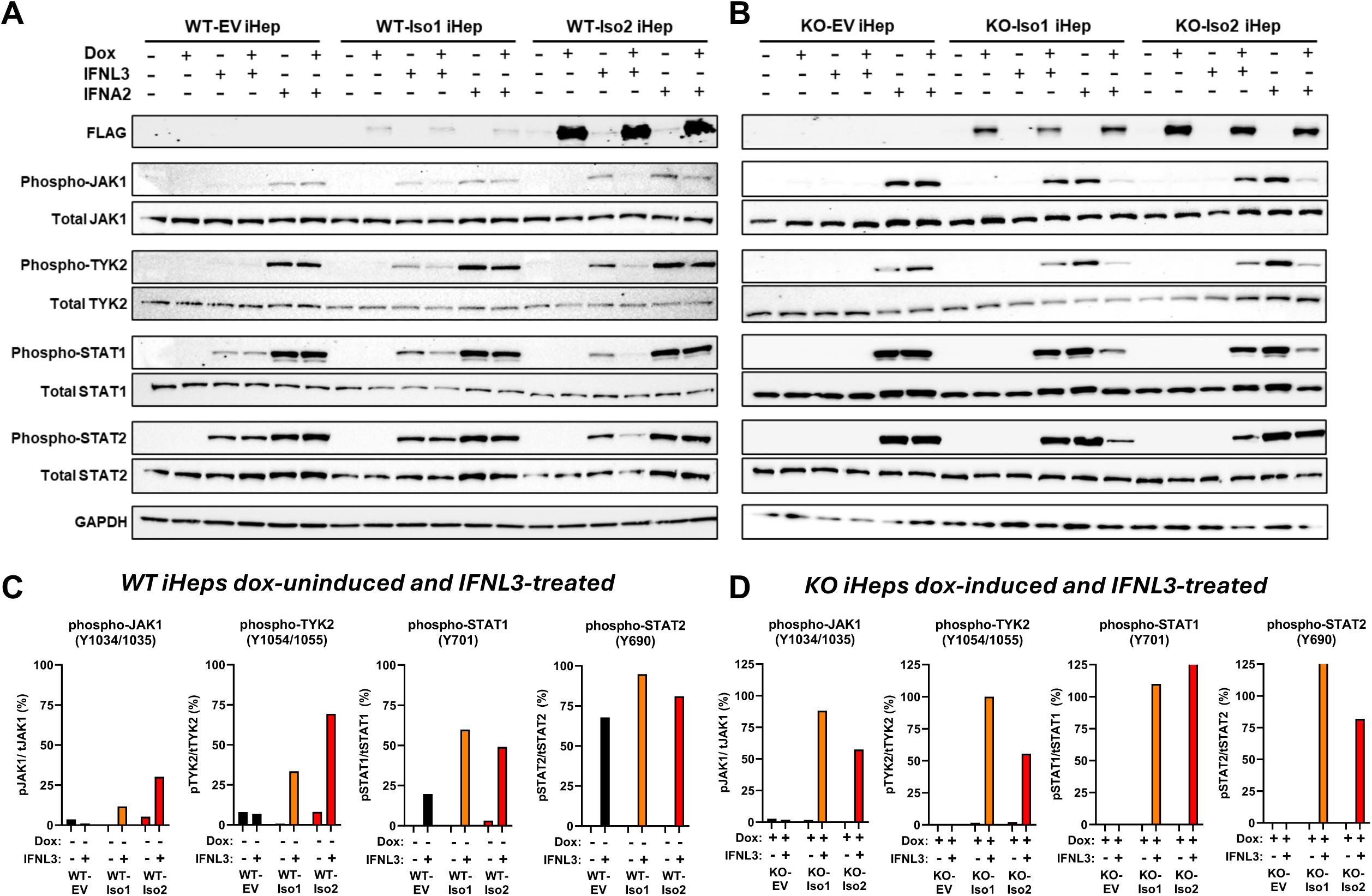
IFNLR1 variants differentially activate JAK-STAT signaling mediators after IFNL3 stimulation. Western blot analysis of whole cell lysates from (**A**) WT iHeps and (**B**) *IFNLR1*-KO iHeps +/-dox-induced for 24h then treated +/-IFNL3 or IFNA2 for 15min. GAPDH served as an indicator of equivalent protein loading per lane. The ratio of phosphorylated to total protein for (**C**) dox-uninduced WT iHeps and (**D**) dox-induced *IFNLR1*-KO iHeps +/-IFNL3 is shown as a percentage, based on integrated band intensity determined in ImageJ.

### Differential susceptibility of IFNLR1 variants to JAK1 and TYK2 inhibition

IFN-modulating therapies are increasingly used as therapeutics^38,66,76^, including inhibitors specific for individual JAK proteins. Given the observed differences in how IFNLR1 variants support JAK1 and TYK2 phosphorylation and subsequent signal transduction, we next evaluated if signaling through FLAG-Iso1 and FLAG-Iso2 were differentially influenced by the clinically utilized inhibitors upadacitinib (an ATP-competitive JAK1 inhibitor) and deucravacitinib (an allosteric TYK2 inhibitor). We compared relative inhibition not only between the variants, but also between WT- and KO-iHeps to evaluate how protein from endogenous *IFNLR1* influenced signaling by the co-expressed FLAG-tagged variants. We quantified the expression of a representative antiviral ISG (*MX1*), whose induction is supported by both FLAG-Iso1 and -Iso2, and a proinflammatory ISG (*CXCL10*), whose induction is only supported by FLAG-Iso1^33–35^. We emphasized evaluation of dox-uninduced conditions for WT-iHeps, to mitigate the risk of off-target effects, but also assayed dox-induced WT-iHeps to enable comparison to KO-iHeps, which require dox-induction to have an observable phenotype from FLAG-tagged variant expression^33–35^.

While WT-Iso1 and KO-Iso1 iHeps had comparable susceptibility to JAK1 inhibition of *MX1* induction, WT-Iso1 iHeps had markedly reduced susceptibility to TYK2 inhibition (**Fig. 4A**). Both lines could be fully inhibited by JAK1 inhibition, but neither line exhibited complete abrogation of *MX1* induction by TYK2 inhibition (**Fig. 4A**). These data suggest noncanonical variants derived from endogenous *IFNLR1* may reduce the TYK2-dependence of IFNL signaling for antiviral ISG induction. Although deucravacitinib has a lower IC_50_ (0.2nM) than upadacitinib (47nM)^77^, FLAG-Iso1 was notably less susceptible to TYK2 than to JAK1 inhibition, particularly in WT-iHeps (**Fig. 4A**). These findings are consistent with prior observations *in vitro* and *in vivo* that IFNL signaling can be TYK2 independent^54–56^.

**Figure 4.**
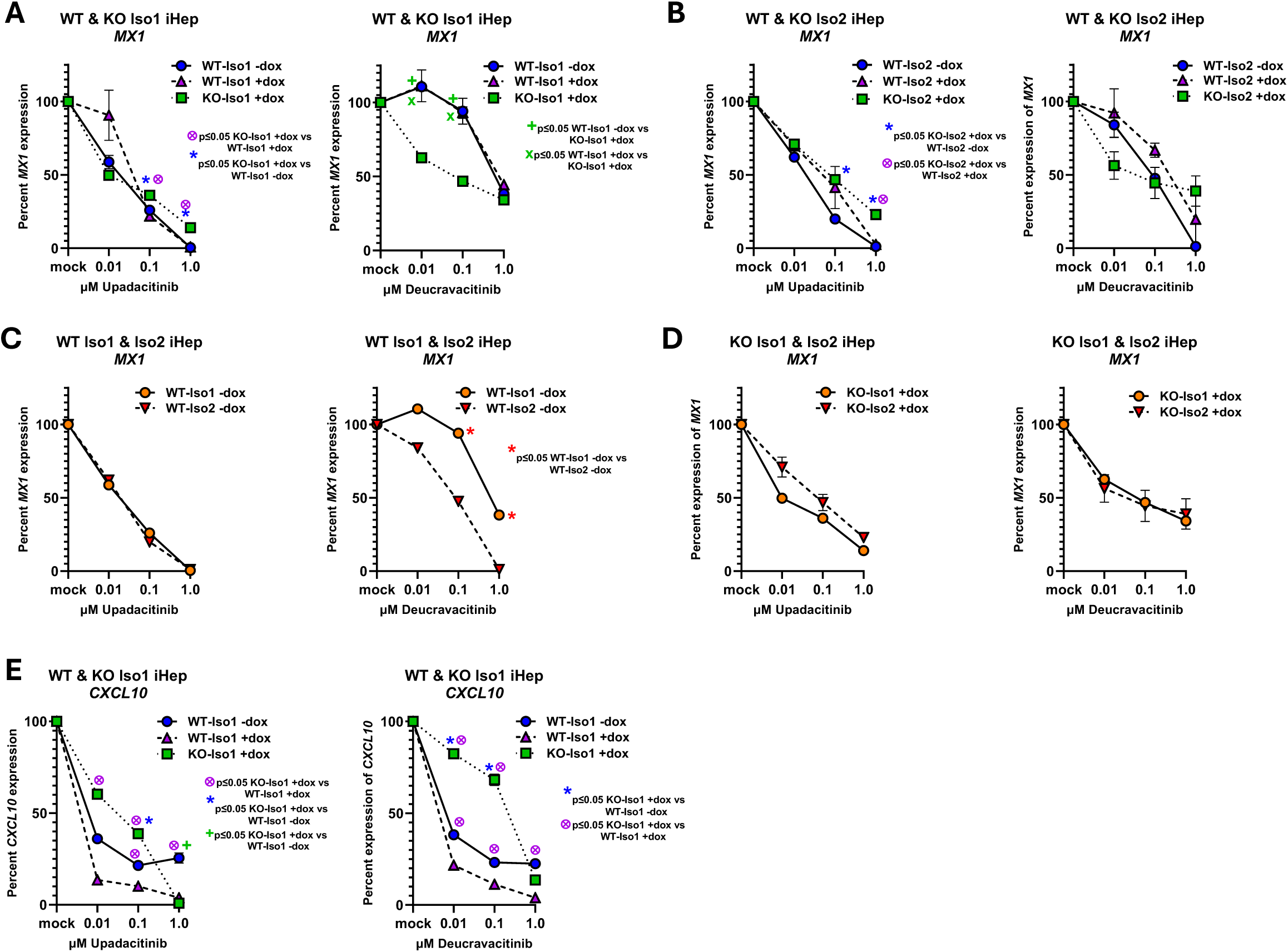
Differential susceptibility of IFNLR1 variants 1 and 2 to JAK1 and TYK2 inhibition. Shown are *MX1* and *CXCL10* expression in iHeps pre-treated with JAK1-upadacitinib; IC_50_ 47nM) or TYK2-(deucravacitinib; IC_50_ 0.2nM) inhibitors prior to IFNL3 stimulation (100ng/ml, 24h) relative to mock-treated cells. (**A**) Comparison of *MX1* expression in WT-Iso1 (+/-dox) and KO-Iso1 (+dox) iHep lines and (**B**) WT-Iso2 (+/-dox) and KO-Iso2 (+dox) iHep lines. (**C**) Comparison of *MX1* expression in WT-Iso1 and WT-Iso2 iHep lines (-dox) and (**D**) KO-Iso1 and KO-Iso2 iHep lines (+dox). (**E**) Comparison of *CXCL10* expression in WT-Iso1 (+/-dox) and KO-Iso1 (+dox) iHeps. Biological replicates were assayed in technical duplicate and mean ± SEM is shown relative to *GAPDH*. Results are representative of two independent experiments. Percent gene expression was calculated relative to respective mock treated samples. *, +, and _ indicate p≤ 0.05 by Student’s t-test.

Similar to WT- and KO-Iso1 iHeps, WT- and KO-Iso2 iHeps were less susceptible to TYK2 than to JAK1 inhibition; however, between cell lines, TYK2 and JAK1 inhibitor susceptibilities were more comparable (**Fig. 4B**). When we compared *MX1* expression in dox-uninduced WT-iHeps expressing each variant, we observed similar susceptibility to JAK1 inhibition, but a notable reduction in susceptibility to TYK2 inhibition in WT-Iso1 relative to WT-Iso2 iHeps (**Fig. 4C**), with full inhibition of *MX1* observed in deucravacitinib-treated WT-Iso2 iHeps. Reduced susceptibility to TYK2 inhibition was similarly observed in HEK293T cells expressing FLAG-Iso1 relative to FLAG-Iso2 (**Supplemental Figure 4**), however no difference in TYK2 susceptibility was observed in the corresponding variant-expressing, dox-induced KO-iHeps (**Fig. 4D**). When we evaluated *CXCL10* expression in iHeps expressing FLAG-Iso1, we found WT-Iso1 iHeps unexpectedly had enhanced susceptibility to both JAK1 and TYK2 inhibition of *CXCL10* expression relative to KO-Iso1 iHeps (**Fig. 4E**), implying endogenous noncanonical variants may increase the dependence on TYK2 and JAK1 for proinflammatory ISG induction, an opposite pattern to what we observed for antiviral ISGs (**Fig. 4A**). These data identify a complex relationship between co-expressed canonical and noncanonical IFNLR1 variants and the TYK2-dependence of IFNL signaling to support induction of antiviral and proinflammatory ISGs.

### Influence of IFNLR1 variants on IFNL-induced gene expression

To further explore how IFNLR1 variants influence the depth and breadth of IFNL-induced genes, we performed RNA-Seq of variant-expressing iHeps in WT (dox-uninduced) and KO (dox-induced) backgrounds (DEGs are reported in **Supplemental File 1**). Consistent with our prior published results that examined select antiviral and proinflammatory ISGs^33–35^, both variants supported IFNL-induced ISG expression, however WT-Iso1 iHeps induced a greater number of ISGs represented within interferon and cytokine pathways than WT-Iso2 iHeps (**Fig. 5A-B**), with no notable differences in down-regulated pathways (**Supplemental Fig. 5A-B**). Consistently, dox-induced KO-Iso1 iHeps had representation of interferon and cytokine pathways that were not detected in KO-Iso2 iHeps using the statistical significance criteria for pathway analysis (**Supplemental Fig. 5C-D**). Comparison of relative expression of the DEGs identified in the top-ranked pathway of IFNL3-treated WT-Iso1 iHeps (interferon alpha/beta signaling) revealed that most ISGs followed an expression pattern similar to *MX1*, wherein increased IFNL3-induced expression was observed in both FLAG-Iso1 and FLAG-Iso2 expressing WT and KO-iHeps relative to the respective EV controls, with maximal expression supported by FLAG-Iso1 relative to FLAG-Iso2 (**Fig. 6A-B**).

**Figure 5:**
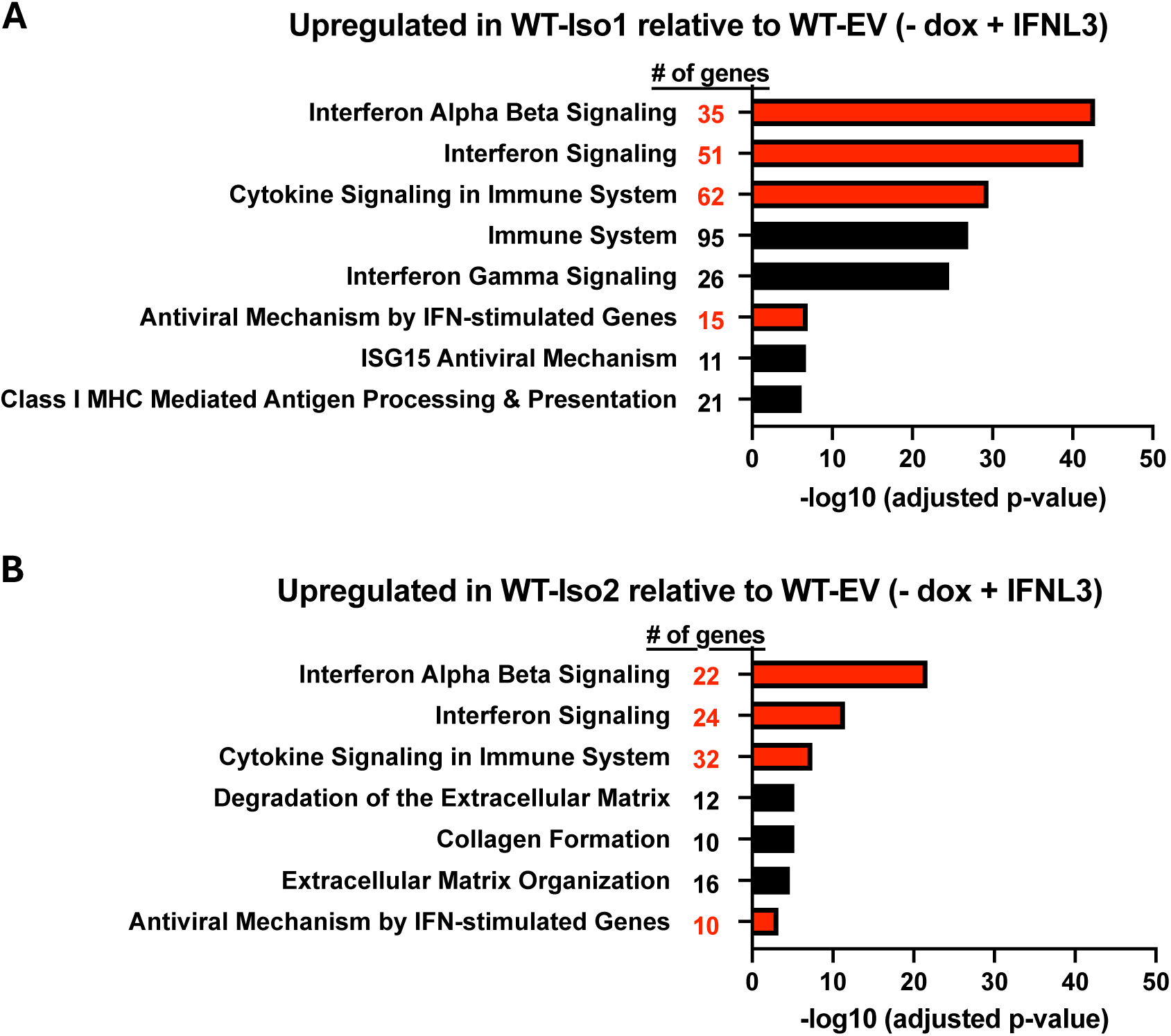
IFNLR1 variant 1 supports broader induction of ISGs than variant 2. Top pathways up-regulated by IFNL3 treatment in dox-uninduced WT-iHeps expressing FLAG-Iso1 (**A**) or FLAG-Iso2 (**B**) relative to similarly treated WT-EV iHeps. Pathways with at least 10 genes identified in the dataset represented within the indicated pathway are shown. Pathways identified in both datasets are indicated in red, with the number of genes from the dataset represented within each individual pathway indicated.

**Figure 6:**
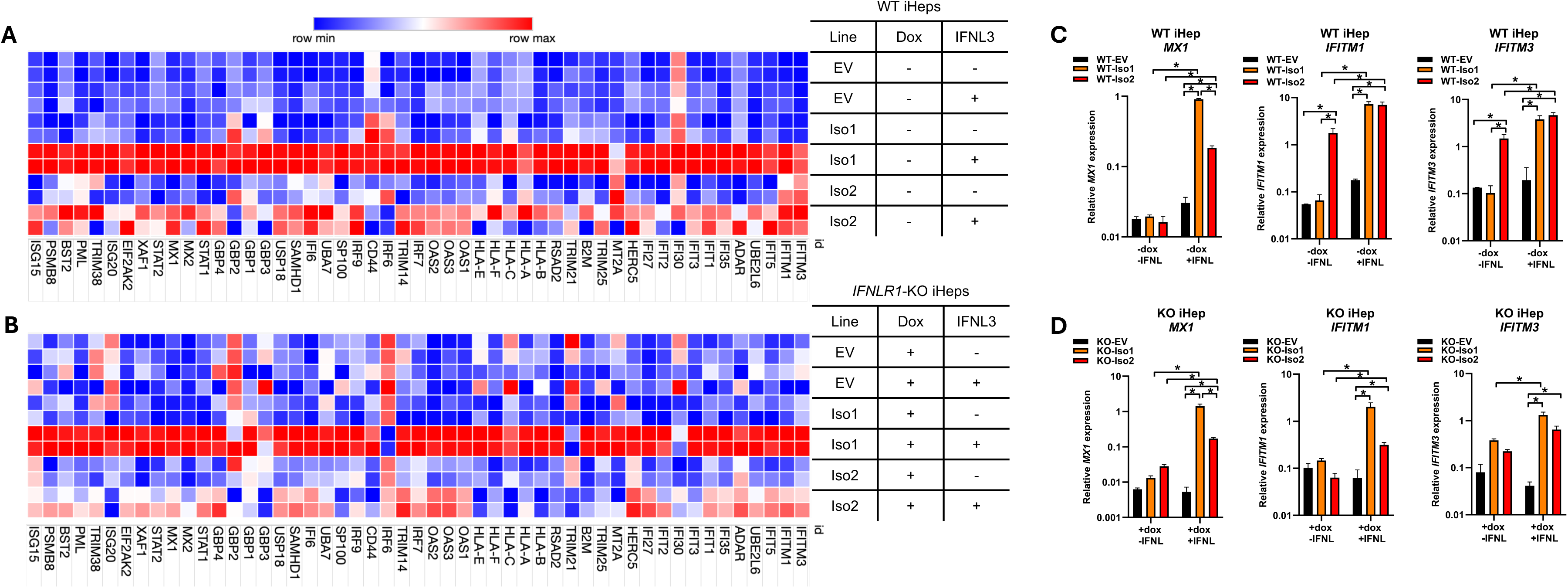
IFNLR1 variants differentially influence constitutive and IFNL3-induced ISG expression. Heat map of RNA-seq data showing column-normalized expression for the 35 IFNL3-induced DEGs identified in the “Interferon alpha beta signaling” pathway in WT-iHeps expressing FLAG-Iso1 relative to EV (-dox, + IFNL3) shown in (**A**) WT and (**B**) KO iHep samples with dox and IFNL3 treatment as indicated. Rows depict individual biological duplicate samples clustered by cell line and treatment; red indicates higher expression and blue represents lower expression (**C-D**) Relative expression of select genes (*MX1*, *IFITM1*, *IFITM3*) evaluated by qRT-PCR relative to *GAPDH* in WT (**C**) and KO (**D**) iHeps showed correlation with RNA-seq data. Data are representative of two biological replicates assayed in technical duplicates relative to *GAPDH*. Mean ± SEM, *p≤0.05 by Student’s t-test.

Interestingly, several ISGs exhibited differential constitutive expression in the absence of IFNL3 treatment, for example *IFITM1* and *IFITM3*, which were expressed in untreated WT-Iso2 iHeps but not in untreated WT-EV or WT-Iso1 iHeps (**Fig. 6A, C**). This pattern was not observed in KO-Iso2 iHeps (**Fig. 6B, D**), however both of these ISGs were IFNL-inducible in all variant-expressing lines. These data are intriguing as *IFITM1* and *IFITM3* exhibit homeostatic expression in certain cellular contexts^78–81^ and IFNLR1 influences homeostatic expression of select ISGs in the intestinal epithelium^6,36,82^. As such, these data suggest noncanonical IFNLR1 variants could influence expression levels of both constitutively expressed and IFNL-induced ISGs.

## Discussion

Using FLAG-tagged native IFNLR1 variants, we evaluated the signaling capacity of full-length canonical IFNLR1 and a noncanonical variant that has identical extracellular and transmembrane domains but lacks part of the cytoplasmic Box1 and Box2 JAK1-interacting motifs. We identified that both variants colocalize with IL10RB after IFNL3 treatment and support the phosphorylation of JAK1, TYK2, STAT1, and STAT2, which leads to induction of ISGs. However, despite comparable binding to IL10RB (Fig. 2A-B) and retention of the cytoplasmic domain tyrosines that enable STAT1/2 binding and phosphorylation^31,37,74,75^, the kinetics and rate of receptor internalization (Fig. 2C-D), nature of engagement with JAK-STAT signaling molecules (Fig. 3-4), and breadth and depth of downstream induced ISGs differed between variants (Fig. 5-6). Our data suggest the missing Box1 and Box2 motifs still permit JAK1 binding and phosphorylation, however the cytoplasmic domain truncation may weaken or destabilize JAK1 and/or pJAK1 binding relative to variant 1, resulting in reduced STAT1/2 phosphorylation (modeled in Fig. 7A). Differential engagement with the proximal signaling kinase (JAK1) by IFNLR1 variant 2 could thus function as a mechanism to enable cells to titer expression of IFNL-induced antiviral ISGs while also limiting induction of proinflammatory ISGs (Fig. 5-6 and ^34,35^). Intriguingly, we also identified that variants produced from endogenous *IFNLR1* influence the TYK2-dependence of IFNL signaling that leads to induction of antiviral vs. proinflammatory ISGs (Fig. 4).

**Figure 7:**
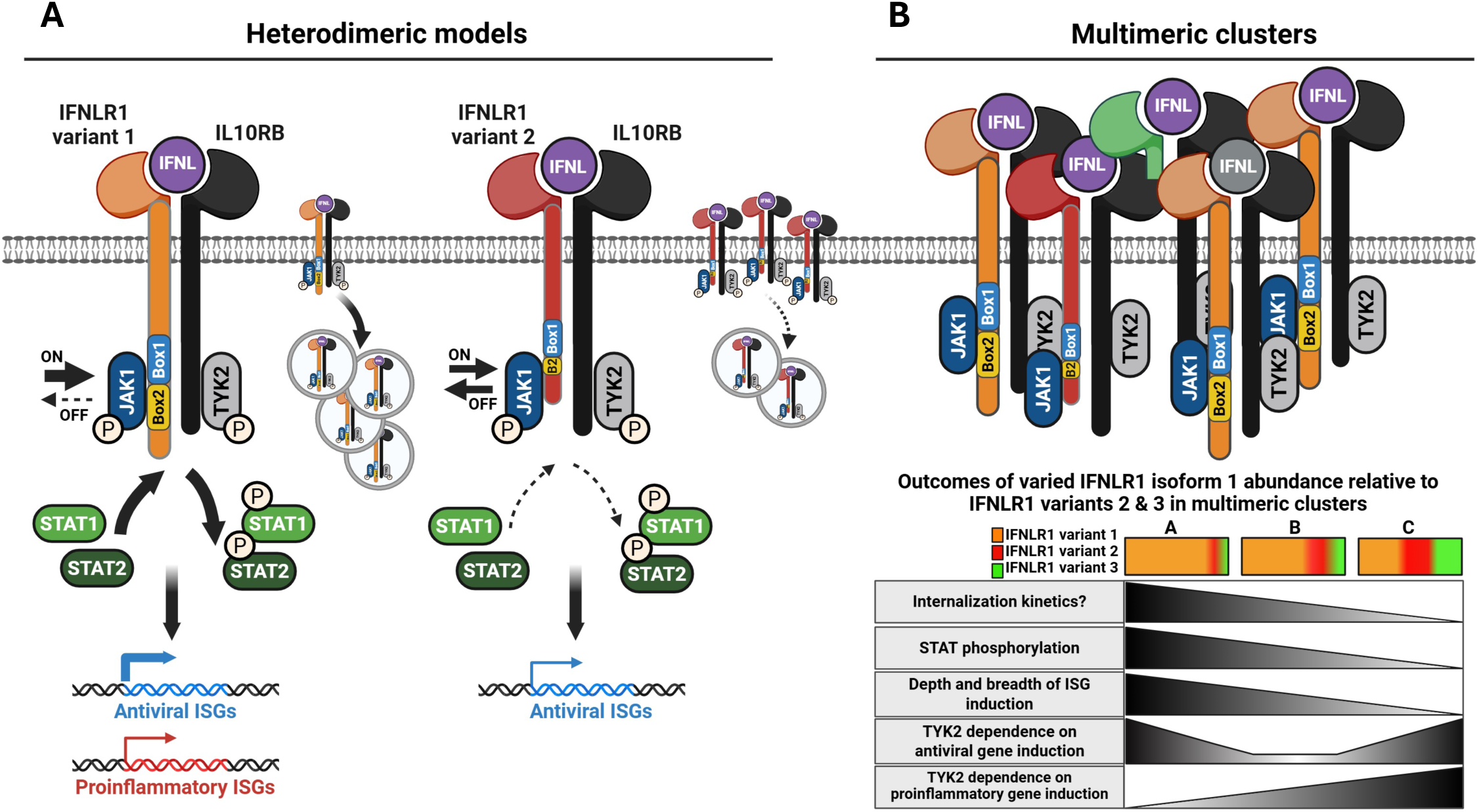
Model of IFNL signaling supported by IFNLR1 variant 1 and 2. (**A**) Model depicting the mechanisms of distinct signaling outcomes imparted by ternary complexes composed of IFNL3, IL10RB and either IFNLR1 variant 1 or 2. Greater arrow width indicates higher association and/or phosphorylation (JAK1, STAT1, STAT2), greater internalization of receptor complexes, or higher induction of gene expression. IFNLR1 variant 1 containing heterodimers are depicted to have greater stability of JAK1 and/or pJAK1 binding and to be more prone to internalization than IFNLR1 variant 2 containing heterodimers. This correlates with variant 1 mediating more efficient phosphorylation of STAT1 and STAT2 and supporting higher expression of antiviral ISGs and *de novo* expression of proinflammatory ISGs compared to variant 2. (**B**) Model depicting assemblage of multimeric clusters containing multiple heterodimers of IFNL3-bound IFNLR1 variants in complex with IL10RB. In this model, proximity of JAK1 molecules bound to the cytoplasmic domains of IFNLR1 variants 1 and 2 could participate in TYK2-independent transphosphorylation. The relative abundance of IFNLR1 variants within multimeric complexes would influence the nature of receptor internalization, STAT phosphorylation, and downstream gene expression. The TYK2 dependence of signaling differs for antiviral vs. proinflammatory ISG expression and has a complex relationship with the relative abundance of IFNLR1 variants. Soluble variant 3, not studied in these experiments, is included in the model for consideration. Created in BioRender. Novotny, L. (2025) https://BioRender.com/24aqcgc.

These data provide potential insight into the observations that IFNL signaling through the IFNLR1/IL10RB heterodimer appears dependent on TYK2 bound to IL10RB^57–59^. In contrast, IFNL signaling was shown to be TYK2-independent in individuals with genetic TYK2 deficiency or those treated with TYK2 inhibitors^36,54–56^. Synthetic IFNLR1 homodimers each bound to JAK1, but not TYK2, propagated IFNL signal independent of TYK2 and IL10RB^57–59^, which suggested JAK1 proteins alone can support IFNL signaling when in proximity and that more complex receptor assemblies other than single IFNLR1/IL10RB heterodimers could form upon ligand exposure. Interestingly, the crystal structure of JAK1 in complex with IFNLR1 suggested a single IFNLR1 protein has theoretical potential to interact with two separate JAK1 molecules^72^. These observations suggest that if aggregation of multiple IFNLR1/IL10RB heterodimers brings the cytoplasmic tails of distinct ternary receptor complexes in proximity, this could provide an opportunity for proximal JAK1-JAK1 molecules to trans-phosphorylate one another and/or for multiple transient interactions of a single JAK1 with multiple IFNLR1 variants, which could obviate or reduce the dependence on TYK2 to propagate signal.

Taken in context with our data, it is intriguing to speculate that IFNLR1 variant 2, IFNL, and IL10RB ternary complexes could colocalize within complex multimeric clusters that also include IFNLR1 variant 1, IFNL, and IL10RB ternary complexes (modeled in Fig. 7B). In this model, the relative abundance of variants that distinctly transduce signal through differential interactions with JAK1 would dictate and diversify the nature of downstream ISG induction (Fig. 7B). As noted above, a less stable interaction of JAK1 with variant 2, or a different conformation of the distal portion of the C-terminal cytoplasmic domain that interacts with STAT proteins, relative to variant 1, could permit mobility or differential exposure of JAK1 within multimeric receptor clusters, bringing it in proximity to other receptor-bound JAK1 molecules. In this model, there would be potential for JAK1-JAK1 transphosphorylation. Considering this model, we can speculate the reduced TYK2 inhibitor susceptibility we observed in WT-relative to KO-iHeps expressing FLAG-Iso1 (Fig. 4A) for antiviral ISG expression (*MX1*) could be explained by variant 2 providing an alternate JAK1 binding site within multimeric clusters that would reduce the need for TYK2 bound to IL10RB to propagate signal. Similarly, the increased JAK1 and TYK2 inhibitor susceptibility of WT-relative to KO-iHeps expressing FLAG-Iso1 for proinflammatory genes (Fig. 4E) suggests noncanonical variants could impair the capacity of variant 1 to transduce signal that leads to induction of proinflammatory genes through a mechanism that is more TYK2-dependent. The differential susceptibility to TYK2 inhibition we observed in WT-but not KO-iHeps expressing FLAG-Iso1 and FLAG-Iso2 (Fig. 4C-D) suggests multiple variants may be required within multimeric signaling clusters to enable disparate downstream signaling outcomes that differ in their dependence on TYK2 bound to IL10RB (Fig. 7B). Our data suggest the TYK2 dependence for IFNLs to induce antiviral ISGs depends in a complex way on the proportion of variant 1 and noncanonical variants, with greater dependence on TYK2 with high or low relative levels of variant 1 and reduced dependence when variants have more even expression (based on data in Figure 4A-D and modeled in Fig. 7B). Finally, our ability to fully inhibit ISG expression with JAK1 inhibitors but not TYK2 inhibitors is consistent with a model wherein JAK1 is required for all avenues of signaling through IFNLR1 in iHeps.

Prior work using synthetic heterodimeric type-I and IFNL receptors showed that differences between IFNAR2 and IFNLR1 in the Box1 and Box2 JAK1-binding motifs accounted for the attenuated STAT1/STAT2 phosphorylation seen in IFNL relative to type-I IFN signaling^59^. Further, differences in signaling between these synthetic heterodimers were not influenced by the distal elements of their respective cytoplasmic domains that harbor the tyrosines responsible for STAT binding and phosphorylation^59^. The differences we observed in the capacity of variants 1 and 2 to transduce signal after promoting JAK1 phosphorylation are consistent with this work, and suggest that the molecular details of JAK1 binding, phosphorylation, and subsequent signal transduction are pivotal for diversifying downstream IFNL signaling outcomes. As JAK proteins are utilized by multiple receptors to propagate signal and JAK binding to Box motifs on receptors can be promiscuous^72,83^, subtle changes in the relative abundance of IFNLR1 variants could thus function to titrate and guide pathway activity (Fig. 7B) and help enable the variability of IFNL signaling that has been observed in different cell types.

Although *IFNLR1* isoform 2 has low transcript expression relative to isoform 1 in primary cells and in our iHep model^34,35^, our data suggest variant 2 may have greater stability on the cell surface both in the presence and absence of IFNL3 (Fig. 2), suggesting protein derived from isoform 2 could be more abundant than would be predicted by transcript levels alone. While variants 1 and 2 both harbor endocytic motifs in their cytoplasmic tails (i.e. YxxΦ), future work will be needed to evaluate whether the missing Box domain elements in variant 2 impair recognition by endocytic adaptor proteins accounting for the observed slower internalization. Additionally, as IFNAR1/2 complexes continue to signal after endocytosis, and internalization can influence the nature of JAK-STAT signaling^49^, it is possible that endosomal signaling by variant 1 could contribute to more sustained JAK-STAT activation relative to variant 2, an important area to explore in future work. Further, whether expression of variant 2 in primary cells could help explain the lack of correlation between IFNLR1 surface expression and cellular responsiveness to IFNL will require further evaluation^74^.

Our findings indicate endogenous IFNLR1 in iHeps is highly sensitive to changes in the relative abundance of its variants, as modest increases in FLAG-Iso1 or FLAG-Iso2, imparted by a leaky tet-on promoter, were sufficient to significantly alter signaling responses when expressed in the presence of endogenous receptor in WT iHeps (Fig. 3-4). In contrast, in the *IFNLR1*-KO background, levels of each FLAG-tagged variant had more modest effects and required dox-induction to have an observable phenotype. These observations suggest that small shifts in the balance of IFNLR1 variant expression could significantly influence receptor function *in vivo*, further highlighting how cells may tightly regulate IFNL signaling outcomes through relative expression of IFNLR1 variants.

We observed a selective influence of IFNLR1 variant expression on constitutive, basal expression of select ISGs that was contingent upon the presence of intact vs. abrogated expression of endogenous *IFNLR1*, most notably in WT-Iso2 cells that had elevated *IFITM1* and *IFITM3* expression in the absence of ligand exposure (Fig. 6). IFNLR1 expression has previously been shown to influence homeostatic ISG expression in the intestinal epithelium^6,36,56,82^ and individuals who are TYK2-deficient have altered expression of select ISGs at baseline^54,56^. Multiple additional studies identified an impact on IFN receptor expression on the basal expression of select ISGs^52,59,84^. As such, relative expression of IFNLR1 variants in IFNL-unexposed cells could contribute to the constitutive balance of ISG expression in cells prior to infection and IFNL exposure. Whether the differences in receptor internalization we observed between variants (Fig. 2) relates mechanistically to the differences we observed in expression of homeostatic ISGs (Fig. 6) will require further investigation.

Strengths of this study include the use of native IFNLR1 protein variants to investigate function, which enabled the interaction of the extracellular domain of IFNLR1 with IFNL ligand and IL10RB. Limitations include the use of protein overexpression to study function, as even subtle changes in protein levels can alter the stoichiometry of JAK-STAT to receptor proteins and influence the nature of downstream signaling^72,83^. To the extent possible, we mitigated the effect of protein overexpression by performing experiments in dox-uninduced conditions when possible, however experiments in *IFNLR1*-KO iHeps required higher protein expression levels to enable study. While not addressed in this work, a third secreted variant of IFNLR1 may also modulate signaling by binding and sequestering IFNL at the cell surface^22,65^ or by interacting with IL10RB and IFNL within complex receptor assemblages (Fig. 7B), an area that will benefit from further study. Additionally, we did not address possible alternative use of JAK2 or other signaling mediators previously implicated in noncanonical IFNL signaling (PI3K, MAPK, NFkB)^27,38,56,85–87^ that could also have contributed to the observed differences between variants. How IFNLR1 variants affect the composition of STAT dimers^31^ and whether they influence cross-talk between pathways that share use of IL10RB (e.g. IL-10, IL-22)^31^ are also important areas of future work. In addition, further studies to target endogenous expression of canonical and noncanonical *IFNLR1* transcripts will help further clarify the role of IFNLR1 variants in IFNL signaling.

In conclusion, our results suggest a model in which IFNLR1 variant 1 in complex with IFNL and IL10RB undergoes efficient internalization and optimizes STAT phosphorylation to support robust induction of antiviral and proinflammatory ISGs. In contrast, variant 2, with its truncated cytoplasmic Box1 and Box2 motifs, supports robust JAK1 and TYK2 phosphorylation but attenuates STAT1/2 phosphorylation leading to a dampened ISG transcriptional response. In this model, we speculate that ternary IFNLR1 variant 2/IFNL/IL10RB complexes could colocalize in proximity to ternary IFNLR1 variant 1/IFNL/IL10RB complexes within multimeric clusters, providing an alternate avenue for JAK1 transphosphorylation that could enable TYK2-independent IFNL signal transduction. In this model, cells could utilize relative expression of IFNLR1 variants to guide expression of antiviral vs. proinflammatory ISGs in order to regulate IFNL responses at mucosal and immune-tolerant surfaces.

## Acknowledgements

We acknowledge Paige Lamprecht-McGinnis and Christiana S. Kappler for assistance with iHep reagents, Josh Monts for technical assistance with ImageStream data acquisition and analysis, and J. Gray Evans for helpful discussions (all at the Medical University of South Carolina).

## Authorship confirmation/contribution statement

Conceptualization, L.A.N., E.G.M.; Methodology, L.A.N., C.M-M., S.A.D, P.T., M.G., E.G.M.; Investigation, L.A.N., C.M-M, M.G., E.G.M.; Writing—Original Draft, L.A.N., E.G.M.; Writing—Review and Editing, L.A.N., C.M-M., S.A.D., P.T., M.G., E.G.M. All authors have read and agreed to the submitted version of the manuscript.

## Funding

This work was supported by funding EGM received from the National Institute of General Medical Sciences (NIGMS: P20GM130457), the National Institute of Diabetes and Digestive and Kidney Diseases (NIDDK: P30DK123704), and the South Carolina Clinical & Translational Research (SCTR) Institute (NCATS: UL1TR001450). MG was supported by the Chan Zuckerberg Initiative and the Silicon Valley Community Fund (Grant#2021-244254). Flow cytometry facilities were supported by the Flow Cytometry and Cell Sorting Shared Resource, Hollings Cancer Center, Medical University of South Carolina (P30 CA138313). Image facilities were supported in part by the Cell & Molecular Imaging Shared Resource, MUSC Cancer Center Support Grant (P30 CA138313), the SC COBRE in Digestive and Liver Diseases (P20 GM130457), the MUSC Digestive Disease Research Cores Center (P30 DK123704,) and the Shared Instrumentation Grants S10 OD018113 and S10 OD028663.

---

**Supplemental Table 1.**
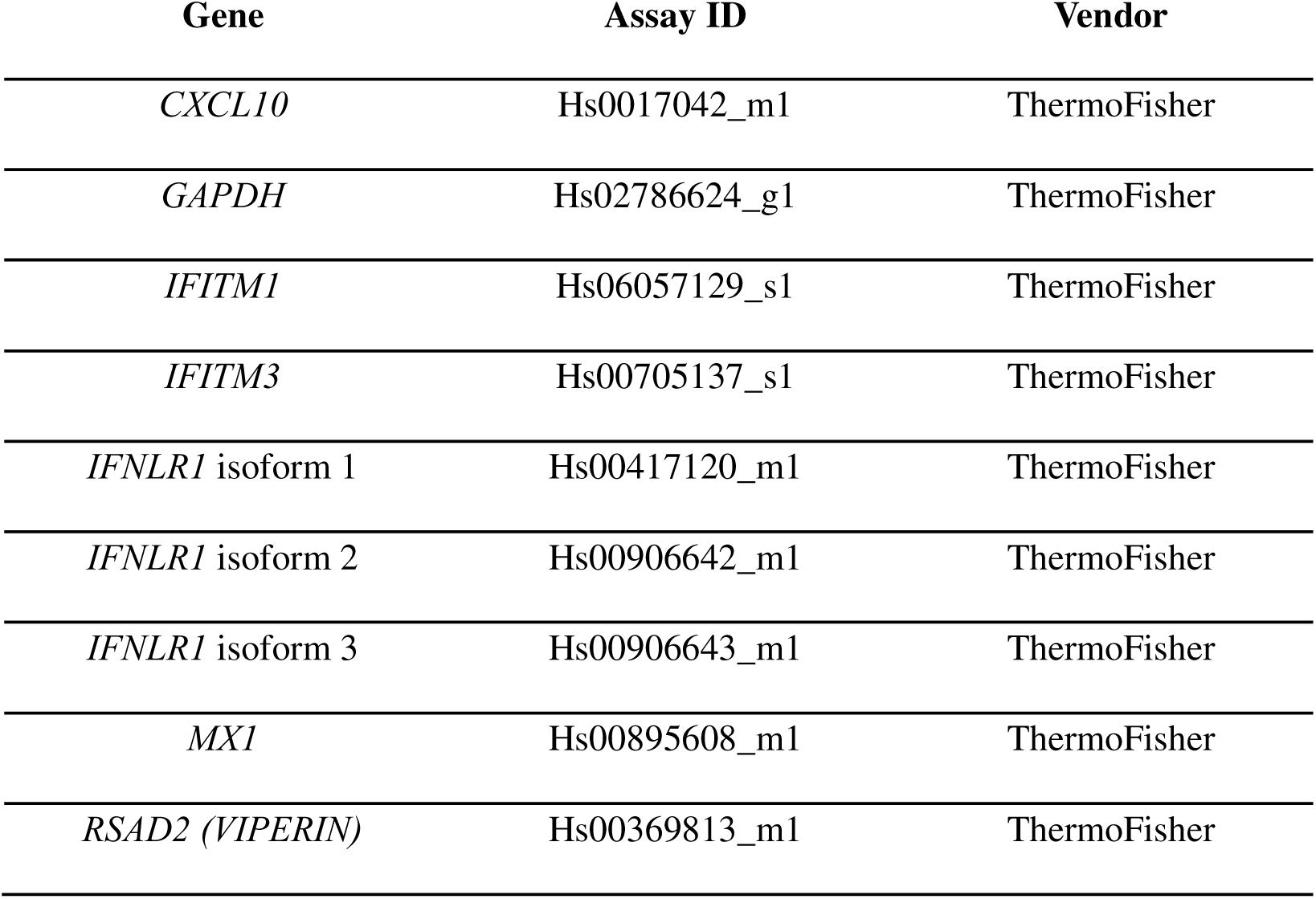
Primer-probe sets used for gene expression analyses.

**Supplemental Figure 1.**
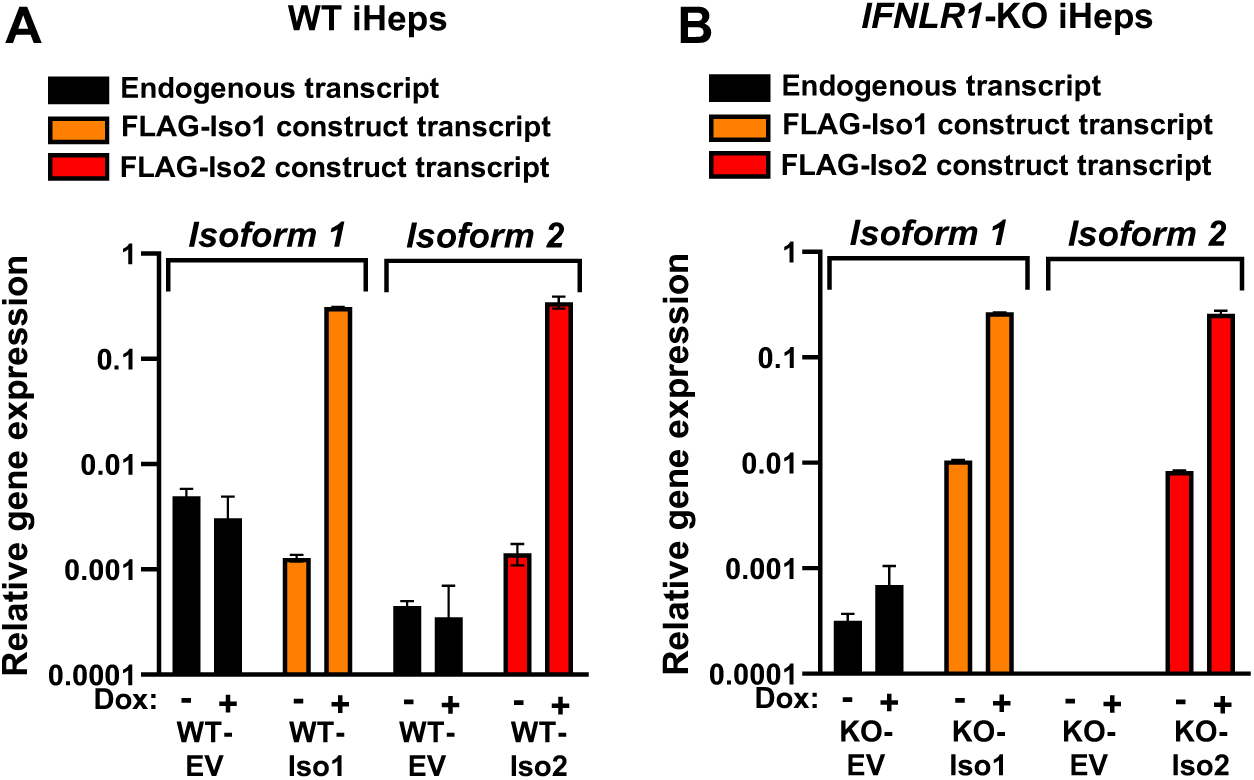
Relative expression of endogenous IFNLR1 and FLAG-Iso1 and -Iso2 transcripts in (A) wild type (WT) and (B) IFNLR1-knock out (KO) iHeps with or without doxycycline induction, determined by qRTPCR. Biological replicates were assayed in technical duplicate and mean ± SEM is shown relative to GAPDH.

**Supplemental Figure 2.**
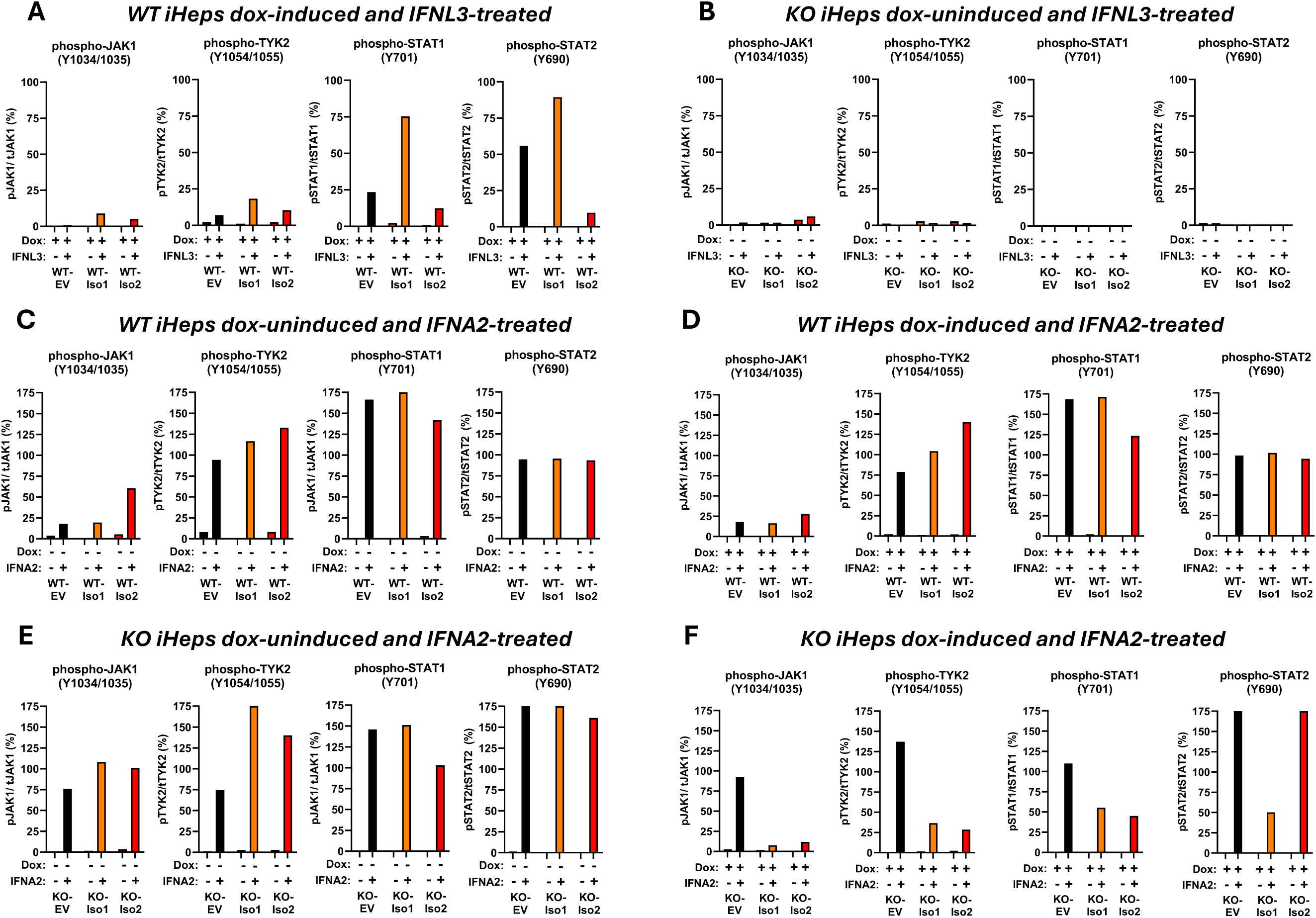
Percentage of phosphorylated to total protein in WT and KO iHep western blots shown in Fig. 4 for (A) dox-induced WT iHeps +/-IFNL3, (B) dox-uninduced IFNLR1-KO iHeps +/-IFNL3, (C) dox-uninduced WT iHeps +/-IFNA2 (D) dox-induced WT iHeps +/-IFNA2, (E) dox-uninduced IFNLR1-KO iHeps +/-IFNA2 and (F) dox-induced IFNLR1-KO iHeps +/-IFNA2 for 15min. Integrated band intensity was determined in ImageJ.

**Supplemental Figure 3.**
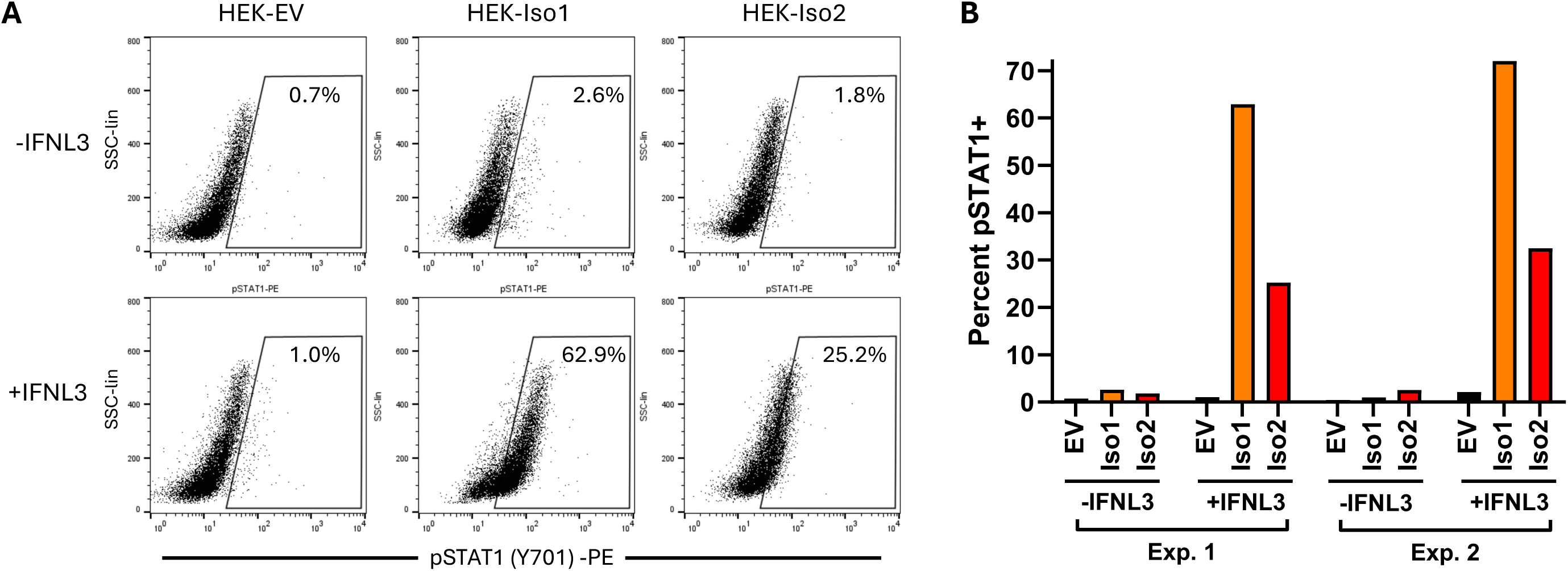
IFNLR1 variant 1 drives greater IFNL3-induced activation of STAT1 compared to variant 2. (A) Representative flow cytometry scatterplots depicting HEK293T cells incubated with or without IFNL3 for 15min then labeled for pSTAT1. The proportion of each population with positive signal is indicated within each scatterplot and summary results from two independent experiments are shown in (B).

**Supplemental Figure 4.**
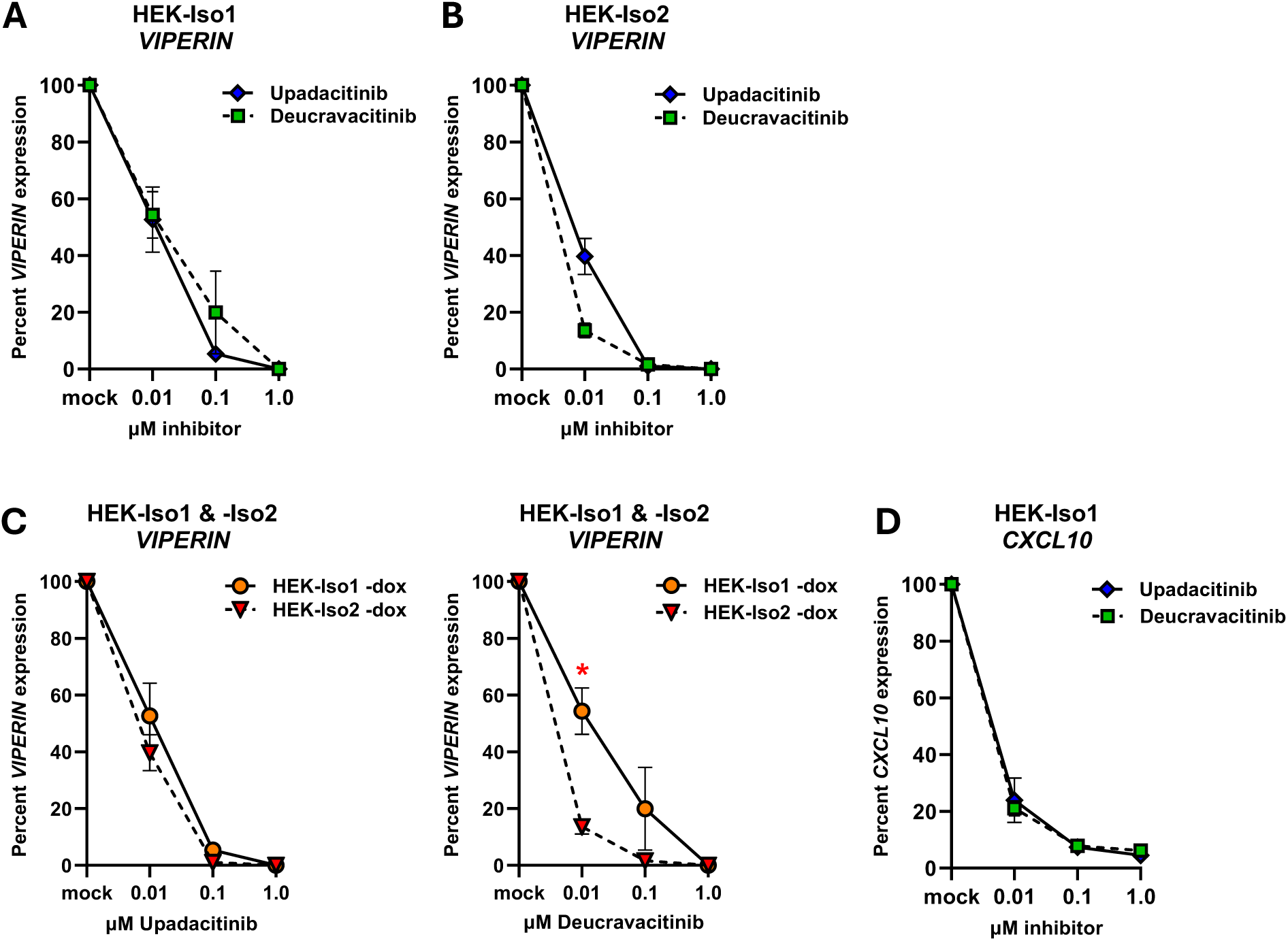
Percent antiviral (VIPERIN) and proinflammatory (CXCL10) gene expression in HEK293T cells pre-treated with JAK1-(Upadacitinib; IC50 47nM) or TYK2-(Deucravacitinib; IC50 0.2nM) inhibitors prior to IFNL3 stimulation. VIPERIN expression in dox-uninduced (A) HEK-Iso1 and (B) HEK-Iso2 cells. (C) Comparison of percent VIPERIN expression between HEK-Iso1 and –Iso2 cell lines. (D) Percent expression of CXCL10 in HEK-Iso1 cells. Biological replicates were assayed in technical duplicate and mean ± SEM is shown relative to GAPDH. Percent gene expression was calculated relative to respective mock treated samples. *p≤ 0.05 by Student’s t-test. Data are representative of two independent experiments.

**Supplemental Figure 5.**
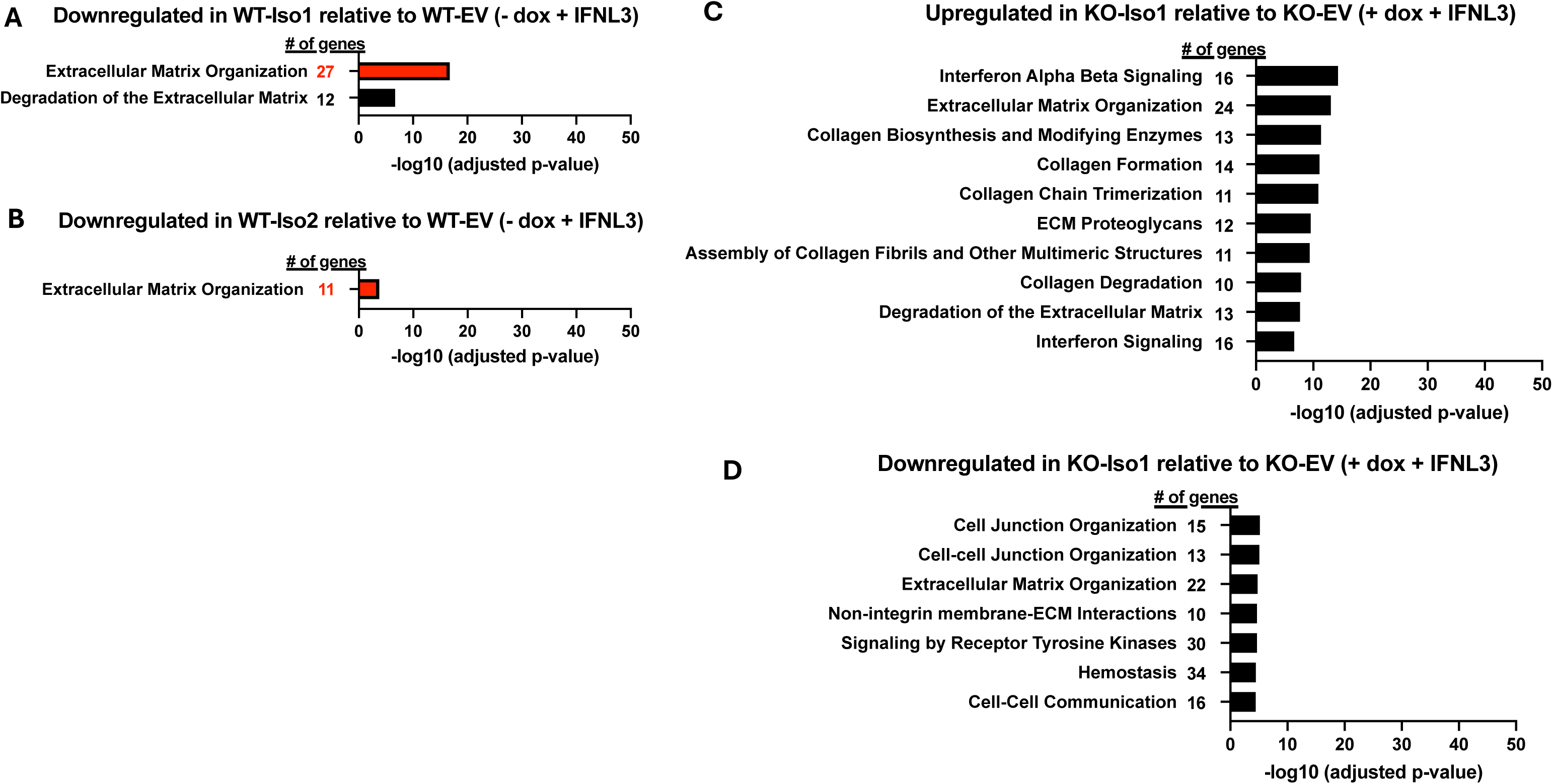
IFNLR1 variants support differential gene expression. Top pathways down-regulated by IFNL3 treatment in dox-uninduced WT-Iso1 (A) or WT-Iso2 (B) iHeps relative to similarly treated WT-EV iHeps. Only pathways with at least 10 dataset genes represented in the pathway are shown. Pathways represented in both datasets are indicated in red, and the number of genes from the dataset represented in each individual pathway are indicated. Top up-(C) and down-regulated (D) pathways in dox-induced, IFNL3-treated KO-Iso1 iHeps relative to similarly treated KO–EV iHeps. No up or down-regulated pathways met the selected significance criteria (see Methods) for KO-Iso2 iHeps.

## Notes

### Competing Interest Statement

The authors have declared no competing interest.

